# The Structural and Biochemical Basis for FER Receptor Kinase Early Signaling Initiation in Arabidopsis

**DOI:** 10.1101/2022.03.30.486374

**Authors:** Yanqiong Kong, Jia Chen, Hong Chen, Yanan Shen, Lifeng Wang, Yujie Yan, Huan Zhou, Heping Zheng, Feng Yu, Zhenhua Ming

## Abstract

Accumulating evidence has indicated that receptor-like kinase (RLK) autophosphorylation and substrate phosphorylation triggered by RLK are early and essential events for RLK function. However, the structural and biochemical basis for these early events is largely unclear. Herein, we used RLK FERONIA (FER) as a model and crystallized its core kinase domain (FER-KD) in the dephosphorylated state. We found that FER-KD adopts an active conformation in its crystal structure. Moreover, FER-KD mutants with reduced or no catalytic activity also adopt an active conformation before phosphorylation. Collectively, these observations suggest that FER employs a phosphorylation-independent active state before ligand-induced phosphorylation and full activation. We further demonstrated that FER is a dual-specificity kinase and that autophosphorylation on Tyr residues lags somewhat behind Ser/Thr phosphorylation. More importantly, Tyr phosphorylation is essential for FER-KD to initiate substrate GRP7 phosphorylation. Our work provides a paradigm to study the mechanisms of the early steps of RLK signaling initiation and highlights its “active” form and Tyr phosphorylation-gated roles in response to signaling stimuli.

## INTRODUCTION

Receptors localized at the plasma membrane (PM) are critical for the adaptation of organisms to different environmental signaling intensity stimuli. The largest family of membrane receptors in animals is the G protein-coupled receptor (GPCR) family, while receptor-like kinases (RLKs) constitute the most prominent family of membrane receptors in plants. In Arabidopsis, there are more than 600 RLKs, making up 2.5% of the protein-encoding genes (**1**). The majority of RLKs have an extracellular ligand-binding domain (ECD), followed by a single transmembrane (TM) helix-containing domain and a relatively well-conserved cytosolic kinase domain (CD) (**2**). The mechanisms of ligand perception of various ECDs have been extensively investigated. For example, leucine-rich repeats (LRR)-RLK brassinosteroid-insensitive 1 (BRI1) senses growth-promoting brassinosteroids (**3**); phytosulfokine receptor (PSKR) recognizes the peptide hormone phytosulfokine (**4**); and root meristem growth factor receptor (RGFR) perceives the RGF peptide (**5, 6**). For a comprehensive review of how RLKs sense various ligands, please refer to Hohmann *et al*. 2017 (**7**). Studies of CDs focus mainly on the identification of key substrates that mediate the downstream regulatory machinery. For example, cognate substrates have been found for Chitin Elicitor Receptor Kinase 1 (CERK1) (**8**), Flagellin Sensing 2 (FLS2) (**9, 10**), BRI1 (**11-14**) and Clavata1 (CLV1) (**15, 16**). Despite advances in understanding ECD-ligand and CD-substrate interactions, the molecular mechanisms of the earliest events of CD activation and subsequent substrate phosphorylation, which leads to the onset of a multitude of cellular processes, remain largely unknown.

CDs are generally composed of three subdomains: the juxtamembrane (JM) domain, core kinase domain (KD) and C-terminal (CT) tail. Apparently, phosphorylation on the CD modulates the activity of the kinase and plays essential roles in signal transduction. Consistent with this notion, phosphorylation of BRI1-associated receptor kinase 1 (BAK1)-CD activates BAK1 and contributes to further phosphorylation of FLS2 (**17**), while phosphorylation of BRI1 enhances its kinase activity and affinity for the coreceptor BAK1 (**14, 18**). RLKs are annotated as serine/threonine (Ser/Thr) kinases, but recent work has shown that tyrosine phosphorylation (pTyr) is also crucial for the activation of RLK-mediated signaling in plants, yet the detailed mechanism of how pTyr could contribute to full kinase activation in these processes is still unknown (**19, 20**). At the same time, sequence analysis revealed that ∼20% of RLKs exist as putative pseudokinases (kinase-defective), lacking one or more residues required for protein phosphorylation (**21**). Notably, some of these putative pseudokinases, which probably function in a phosphorylation-independent manner, have essential roles in distinct biological processes in plants, such as maize atypical receptor kinase (MARK) (**22**), strubbelig (SUB) (**23**), Arabidopsis Crinkly4-related 1/2 (AtCRR1/2) (**24**), guard cell hydrogen peroxide-resistant 1 (GHR1) (**25**), and pangloss 2 (PAN2) (**26**). One possibility is that they act as allosteric regulators or scaffolding molecues to recruit downstream proteins and mediate signal transduction. Consistent with observations on functional pseudokinases, the kinase activity of some RLKs is unnecessary or redundant for certain functions. For example, the “kinase-dead” RLK FER can still rescue its loss-of-function mutant phenotype in some cellular processes (e.g., leaf cell growth and double fertilization) (**27-29**). In sharp contrast, FER kinase activity is pivotal for rapid alkalinization factor 1 (RALF1)-induced root growth inhibition (**30**). Given that a theoretical framework is lacking for deciphering the sequence-activity-function relationship (especially the interplay between), we chose FER as a notable example.

FER belongs to the 17-member *Cr*RLK1L (*Catharantus roseus* RLK1-like) family of RLKs in Arabidopsis (**31**). Recent studies have demonstrated that FER acts as a crucial protein to control many aspects of plant stress responses (e.g., immune responses) and cell growth in a context-specific manner (**32, 33**). In the extracellular environment, FER functions as a receptor for the plant peptide hormone RALFs (**30, 34**). FER is a versatile kinase that inhibits pollen tube growth to affect double fertilization (**35**), suppresses primary root growth by responding to the RALF1 peptide (**30**), and promotes cell growth in root hairs and leaves (**36**). Interestingly, FER activity levels or signaling outputs vary according to the perception of different RALF members (**37**). For example, RALF1 and RALF23 activate FER signaling during root growth, while RALF23 represses FER signaling during immune responses (**30, 34, 38**). These results suggest that FER probably employs its kinase domain to recruit various downstream factors and regulate different biological processes. Given that FER has multiple phosphorylated sites, it is worthwhile to identify the critical sites as the first step to study whether FER could also control its functional outputs via different phosphorylation site patterns. This hypothesis is supported by the results of different mutant lines of FER. *fer-4* is a knockout mutant with a stop codon before the kinase domain (i.e., no kinase domain expression) (**39**), while *fer-5* is a knockdown mutant with deletion of the C-tail of the kinase domain (**39**). Genetic analyses have shown that both *fer-4* and *fer-5* impact the auxin response (**39**), while only *fer-4* has an obvious impact on the ABA response (**40**). A fascinating question is how much kinase activity is needed for multiple FER functions. Accumulating genetic data suggest that defects in certain specific cellular processes (e.g., pollen tube growth regulation) can be rescued by the FER^K565R^ “kinase-dead” mutation (Lys at 565 mutated to Arg; the mutant has no detectable self-phosphorylation level) (**27**), indicating that FER kinase activity may not be needed for this process. However, the FER^K565R^ mutation cannot rescue certain cellular processes, such as RALF1-induced root growth inhibition (context-specific dependence on FERONIA kinase activity) (**41**). These data suggest that FER RLK, unlike BRI1 RLK, may exhibit some unknown general kinase regulation mechanisms of RLKs and/or RLK pseudokinases, such as the mechanism by which RLKs function without complete phosphorylation or catalytic activity. Based on these reasons, we determined the high-resolution crystal structures of FER-KD and its two mutants and identified an unanticipated active state of RLK formed without detectable phosphorylation. Through a combination of mass spectrometry and biochemical analyses, a mechanistic model for the early steps of FER signaling initiation was discovered.

## RESULTS

### The FER core kinase domain adopts an active conformation without being phosphorylated

To study the activation and regulatory mechanism of FER kinase, we expressed, purified and crystallized its core kinase domain (FER-KD, residues 518-816) (**Fig. 1A**). This construct is a truncated form without the JM and CT subdomains, which contains mainly disordered regions, as indicated by the results from HHpred (**42**) and Phyre2 (**43**) analyses. Wild-type FER-KD arrests growth when expressed in *Escherichia coli*, and we hypothesize that the toxicity is related to the kinase activity of FER. To overcome this problem, FER-KD was coexpressed with abscisic acid-insensitive 2 (ABI2), given that ABI2 directly interacted with and dephosphorylated FER (**44**). After purification, FER-KD was crystallized in complex with ADP (FER-KD/ADP), and the complex structure was determined at ∼2 Å resolution (**Table 1**). Consistent with the coexpression of FER-KD with ABI2, careful examination of the crystal structure revealed no phosphorylation on the core kinase domain of FER-KD. To further confirm this notion, we detected the phosphorylation state of the FER-KD proteins in the crystals by using anti-phosphoserine/phosphothreonine (anti-Ser/Thr) antibodies, and no phosphorylation was detected (**Fig. S1A**). The current state of FER-KD likely corresponds to an early step of RLK activation, in which RLK has not been primed by the ligand.

**Figure 1.**
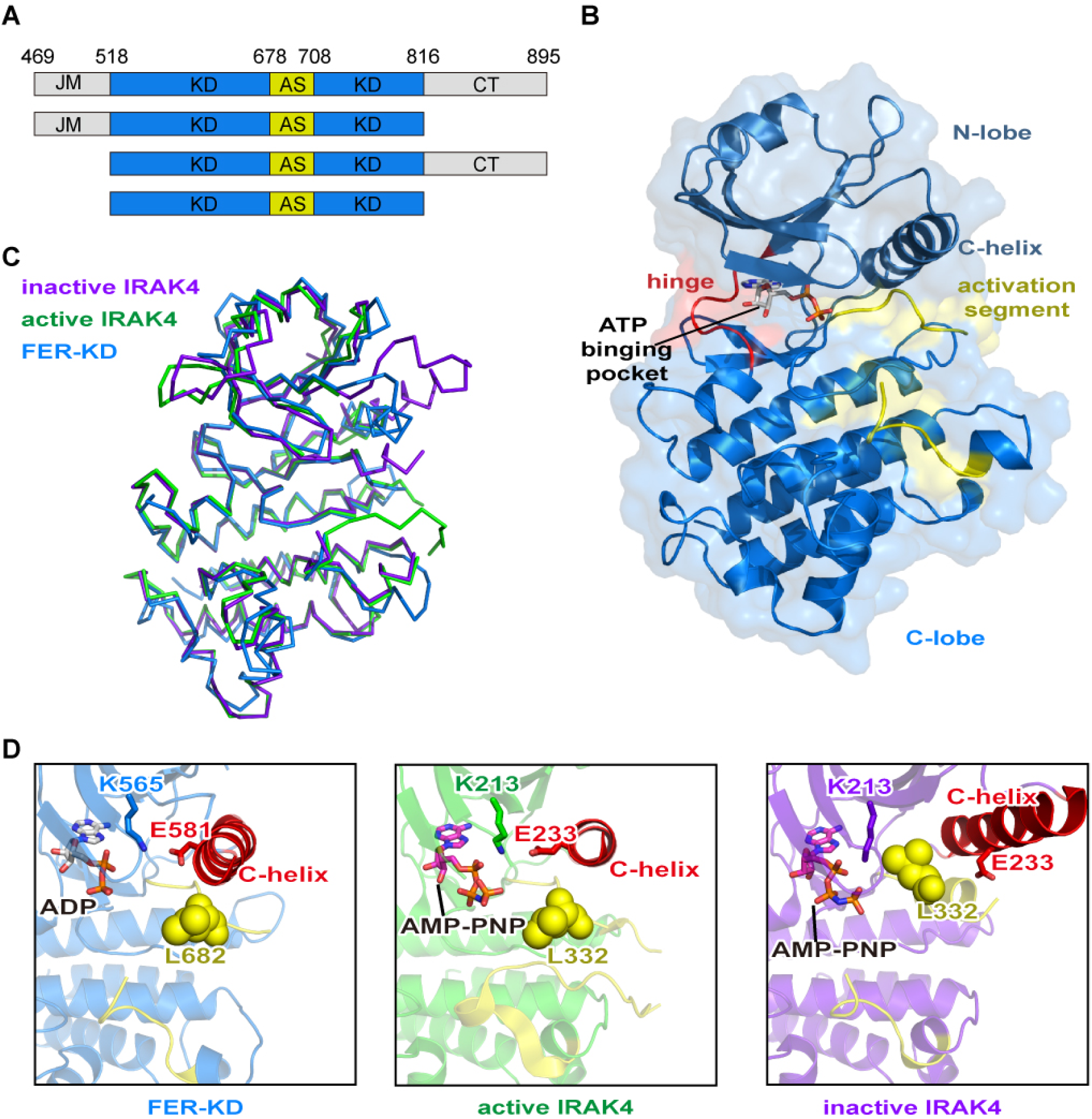
The kinase domain of FER adopts an active conformation. **(A)** Schematic diagram of the FER fragments from the intracellular domain (FER-CD) in our study. JM, juxta-membrane domain; KD, kinase domain; AS, activation segment; CT, C-terminal. Residues located at the domain boundaries are numbered on top of the panel. (B) Overall structure of FER-KD in complex with ADP. The structure of FER-KD, as shown in a cartoon and surface representation, reveals an active conformation. The N-lobe, hinge region, and C-lobe of the kinase are colored dark blue, red and sky blue, respectively. Key structural elements responsible for substrate binding and catalysis (the ATP-binding pocket, activation segment and C-helix) are highlighted. ADP is shown as colored sticks. (C) Structural comparison of FER-KD (dark blue) with active IRAK4 (green, PDB ID 2OID) and inactive IRAK4 (purple, PDB ID 6EGF). Superimposition of FER with active and inactive IRAK4 gives Cα RMSDs of 0.815 Å and 1.104 Å, respectively, supporting the notion that FER is in an active state. All three structures are represented as ribbons. (D) Detailed comparison of the C-helix and activation loop of FER-KD, IRAK4 in the active conformation and IRAK4 in the inactive conformation. The activation segments are colored yellow. The C-terminal region is colored red. The regulatory leucine residue (L682 in FER-KD and L332 in IRAK4) is shown as yellow spheres. The catalytic lysine (K565 in FER-KD and K213 in IRAK4) and its conformation-activating glutamate (E581 in FER-KD and E233 in IRAK4) are shown as colored sticks.

**Table 1.** Data collection and structure refinement statistics.

|  | FER-KD - ADP | FER-KD<sup>S525A</sup>-ADP | FER-KD<sup>K565R</sup>-ADP | FER-KD<sup>K565R</sup>-ATP |
| --- | --- | --- | --- | --- |
| **Data collection** |  |  |  |  |
| *a*, *b*, *c* (Å) | 61.795,83.248,62.322 | 61.67,82.95,61.94 | 61.805,83.163,62.048 | 61.95,83.33,62.05 |
| $\alpha$, $\beta$, $\gamma$ (°) | 90.0, 93.038, 90.0 | 90.0,93.298,90.0 | 90.0, 93.143,90.0 | 90.0,93.39,90.0 |
| Space group | P 1 21 1 | P 1 21 1 | P 1 21 1 | P 1 21 1 |
| Wavelength (Å) | 0.979183 | 0.979183 | 0.979183 | 0.979183 |
| Resolution (Å) | 62.234 - 1.957<br>(2.148 - 1.957) | 82.95 - 2.23<br>(2.29 - 2.23) | 61.95 - 1.93<br>(2.03 - 1.93) | 29.76 - 1.97<br>(2.02 - 1.97) |
| Total No. of reflections | 106756 | 96087 | 139536 | 298055 |
| No. of unique reflections | 32641 | 30350 | 42862 | 44514 |
| Completeness (%) | 88.2 | 99.5 | 90.2 | 99.8 |
| *CC*<sub>1/2</sub> | 0.994 | 0.982 | 0.997 | 0.998 |
| Average *I*/$\sigma$ (*I*) | 8.300 | 9.000 | 13.000 | 12.100 |
| *R*<sub>merge</sub> (%) | 10.1 | 11.1 | 5.7 | 8.6 |
| **Refinement** |  |  |  |  |
| No. of reflections used | 28644 | 28857 | 42809 | 44474 |
| *R*<sub>work</sub> (%) | 19.71 | 18.08 | 18.66 | 17.05 |
| *R*<sub>free</sub> (%) | 24.7 | 23.46 | 20.74 | 21.30 |
| *R.m.s.d.* bond distance (Å) | 0.011 | 0.008 | 0.009 | 0.012 |
| *R.m.s.d.* bond angle (°) | 1.400 | 1.215 | 1.310 | 1.277 |
| Average B-value (Å<sup>2</sup>) | 50.33 | 38.89 | 48.23 | 48.22 |
| Protein atoms | 49.86 | 38.01 | 47.92 | 47.66 |
| Ligands atoms | 88.94 | 113.06 | 79.83 | 78.84 |
| Ramachandran plot (%) |  |  |  |  |
| Favored | 96.20 | 96.20 | 98.74 | 98.39 |
| Allowed | 3.62 | 3.80 | 1.26 | 1.61 |
| Outliers | 0.18 | 0.00 | 0.00 | 0.00 |
| PDB-ID | 7XDY | 7XDX | 7XDW | 7XDV |

Similar to the structure of most kinases, FER-KD adopts a canonical bilobed fold in which the smaller N-terminal lobe is rich in β strands (N-lobe, residues 518-611) and the larger C-terminal lobe contains mostly α helices (C-lobe, residues 618-816). The two lobes are connected by a six-residue hinge (residues 612-617), generating a significant cleft where the ATP substrate is located (**Fig. 1B**). Strikingly, the activation segment (residues 678-708) of FER-KD contains an unmodeled 16-residue middle portion that cannot be observed in the FER-KD/ADP crystal structure, suggesting the high flexibility and disordered nature of this segment. The protrusion of the activation segment out from the core structure and the explosion of the ATP and substrate peptide binding pocket suggests that the observed FER-KD structure adopts an active conformation. The abovementioned hypothesis is further supported by the following three pieces of evidence. The first was derived from a structural similarity search using the DALI program (**45**). The overall structure of FER-KD closely resembles the active fold of many kinases identified but not their inactive fold. Specifically, superimposition of FER-KD with active and inactive interleukin-1 receptor-associated kinase 4 (IRAK4) gives Cα RMSDs of 0.815 Å and 1.104 Å, respectively (**Fig. 1C**) (**46**). Structural comparison of FER-KD with IRAK4 reveals that L682 in FER-KD, which should have restricted kinase activation by inserting the activation loop into the ATP binding site, is in an active conformation. Release of the activation loop from the position between the catalytic residue K565 and the C-helix leads to formation of the K565/E581 ion pair and an active DFGin conformation of the conserved DFG motif (residues 679-681) (**Fig. 1D, Fig. S1B**). Whether the DFGin or DFGout conformation is adopted is determined by the relative position of the Phe residue and C-helix in the DFG motif (**47**). The second piece of evidence was derived from the appropriate formation of the regulatory spine, which corresponds to the active conformation of the kinase (**Fig. S1C**). Taylor and coworkers defined the regulatory spine, which consists of HRD-His (H659 in FER-KD), DFG Phe (F680), C-helix Glu+4 (L585), and a residue in the loop preceding strand β4 (L596) (**48**). Typically, the regulatory spine, which plays important structural roles in the active conformation, is broken by the outward facing αC helix in the inactive conformation of the kinase. In our structure, the regulatory spine is formed in an active conformation (**Fig. S1C**). The third piece of evidence is a nomenclature system to cluster kinase conformations based on the location of the DFG-Phe side chain and the direction of the activation loop (**47**). Our analysis clearly identified the FER-KD structure to be in a “BLAminus” conformation in “DFGin” (residues 679-681) (**Table S1**-**S2**), which is the most common active kinase state in the PDB. Collectively, these analyses reveal that the core kinase domain of FER folds into an active conformation without being phosphorylated, a phenomenon also observed in another animal kinase (**47**). However, it differs from the typical active and phosphorylated BRI1 and BAK1 conformations (**17, 49**). Therefore, this state in the activated conformation but without phosphorylation may herald the emergence of a new mechanism.

### Residues in the ATP-binding pocket are essential for FER catalytic activity

Given that ATP binding is a requisite for phosphate transfer of the kinase, we investigated the ATP-binding pocket of FER-KD in detail. The electron densities for the FER-KD structure clearly show the presence of a molecule of ADP in the ATP-binding site, which is sandwiched between the N- and C-lobes (**Fig. 2A**). ADP adopts a functional conformation with its adenine group hydrogen bonding to the M613 backbone amine and the D611 backbone carbonyl group in the hinge. A magnesium ion bonds with N666 and D679 of the DFG motif (residues 679-681) and with the α- and β-phosphates of the nucleotide. In addition, hydrogen bonds between ADP and E620, N666, D679 and G546 of the protein are likely used to strengthen nucleotide binding (**Fig. 2A**). The C-helix is in an active conformation, allowing E581 and K565 to form a conserved and catalytically important salt bridge. Similar to IRAK4, the conserved gatekeeper position of FER-KD is occupied by the conserved tyrosine Y610. Y610 is oriented toward the conserved E581 and forms a hydrogen bond (**Fig. 2A**), which determines the size of the ‘back pocket’. D661 is considered to be responsible for catalyzing the transfer of phosphate groups. Consistent with the abovementioned structural observations, conservation analysis of the FER subfamily revealed that key residues of the ATP-binding pocket (K565, E581, Y610, D661, K663, N666 and D679) are extremely well conserved in the FER subfamily (**Fig. S2**). To verify whether these pocket residues are required for kinase activity, we constructed several mutants and measured their ATPase activity (see Methods). The results showed that these residues participating in ATP binding were essential for subsequent catalysis by the kinase (**Fig. 2F, Fig. S3A**). In contrast, the mutation of a residue outside the ATP-binding pocket, S525A, only partially affected catalytic activity (**Fig. 2F**).

**Figure 2.**
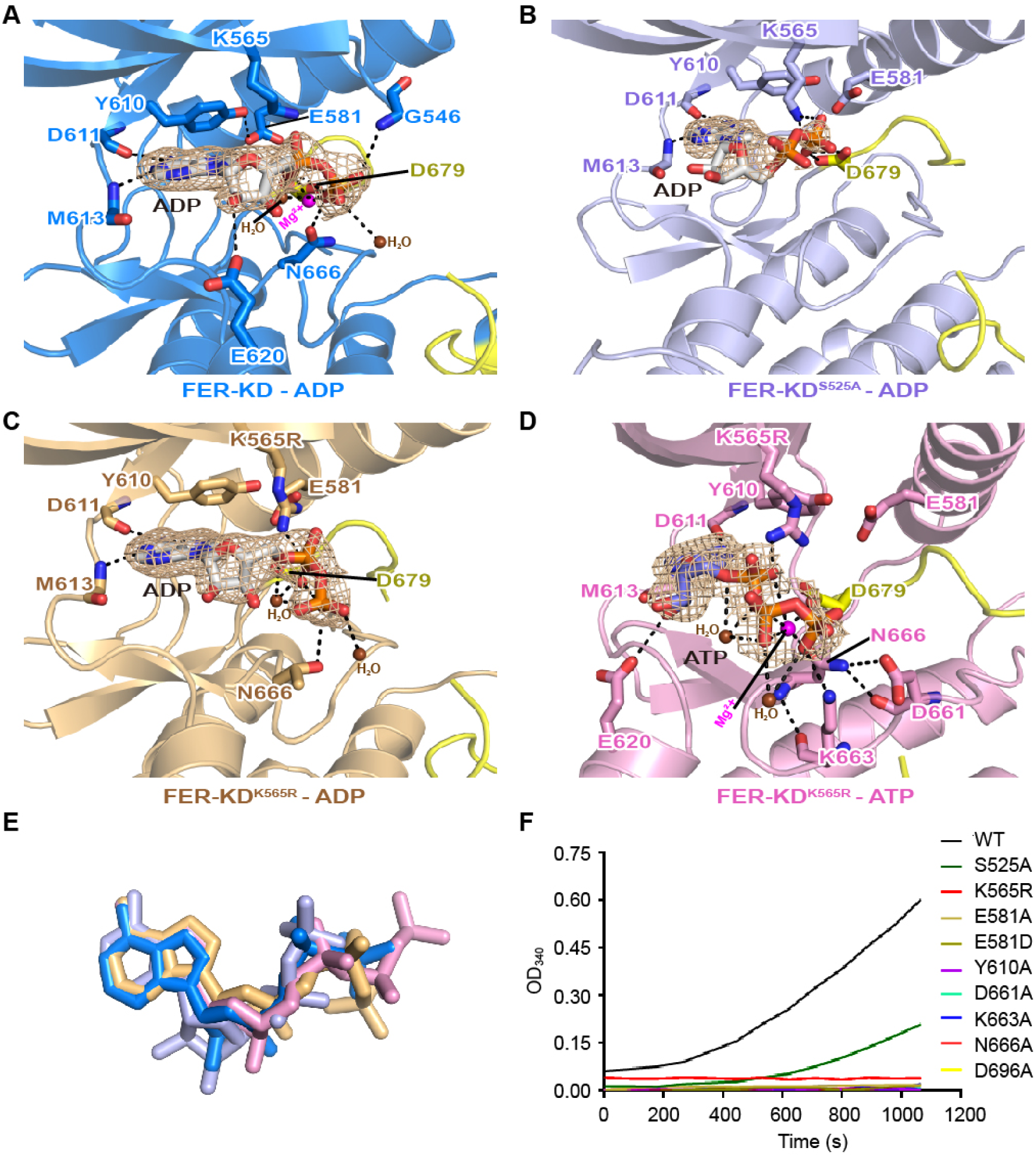
Nucleotides bind in the ATP-binding pocket of FER. **(A)** Close-up view of the ATP-binding site of FER-KD (dark blue), showing the hydrogen bond contacts (black dotted lines) made with ADP, Mg^2+^ and water molecules. The 2*Fo*-*Fc* electron density map, contoured at 1.6σ, is shown in wheat mesh around the ADP molecule. The magnesium ion (magenta) and two water molecules (brown) are highlighted as spheres, whereas the ADP molecule and key contact residues are represented as colored sticks. (**B**), (**C**) and (**D**) Close-up views of the FER-KD^S525A^-ADP (light blue), FER-KD^K565R^-ADP (light orange), and FER-KD^K565R^-ATP (pink) complexes, respectively, in the same orientation and color scheme as (**A**). (**E**) Comparison of the nucleotide orientations in different complex structures. The ADP in structure (**A**) is dark blue, the ADP in structure (**B**) is light blue, the ADP in structure (**C**) is light orange and the ATP in structure (**D**) is pink. The mutation of K565 to R changes the placement of nucleotides. (**F**) ATP consumption activities of FER-KD variants. During the reaction, the phosphorylation process consumes one molecule of NADH for each molecule of ADP produced so that the decrease in the light absorption of NADH at 340 nm (ε340 = 6,220 cm-1 M-1) can be monitored by using a UV spectrophotometer, which then indicates the continuous depletion of ATP. As shown in the figure, the OD340 values of different proteins changed over time. The smaller the difference at the same time, the smaller the ability of the protein to hydrolyze ATP.

### FER mutants that impact the catalytic activity of FER-KD also adopt an active conformation

The K565 site of FER is rather conserved on protein kinases and is generally considered to be associated with ATP binding. Earlier studies have also shown that K565 is an essential catalytic residue, and substitution of this residue by arginine (R) in the FER kinase domain (mFER-KD) causes a dramatic decrease in kinase activity (**41**). By structure and ATPase activity measurements, we also confirmed the essentiality of K565 in ATP binding and subsequent FER activation/autophosphorylation (**Fig. 2F**). We therefore used K565R as an entry point to confirm our previous conclusion that phosphorylation is not obligatory for maintaining the active form of FER. For a more comprehensive view of FER activation, we also chose another FER kinase mutant, S525A, with partially reduced activity (**Fig. 2F**), for subsequent FER activation state analysis. Notably, purified FER-KD^S525A^ and FER-KD^K565R^ were confirmed to have no phosphorylation when coexpressed with ABI2 in *Escherichia coli* (**Fig. S1A**).

We successfully purified and crystallized the mutant K565R in complex with ADP/ATP (FER-KD^K565R^-ADP/ATP) and the partially active mutant S525A in complex with ADP (FER-KD^S525A^-ADP). Consistent with our structural analyses of FER-KD, the mutant structures revealed a similar set of residues for nucleotide binding (**Fig. 2B-D**). Notably, these mutations still allow K/R565 to interact strongly with ADP/ATP molecules, using their positively charged side chains to bind to the negatively charged alpha phosphate group through electrostatic interactions (**Fig. 2B-D**). The retained interaction demonstrates that the ATP-binding affinity may not be compromised, while the main reason for the decreased viability of both S525A and K565R mutants must be due to another reason. Since the K565R mutation results in a significant decrease in the rate of the ATPase reaction (**Fig. 2F**), it is suggested that this lysine site plays a vital role in catalyzing ATP hydrolysis. In the FER-KD^K565R^-ADP and FER-KD^K565R^-ATP structures, the K565R mutation altered the conformation of the catalytic residue D679, which affected the position of the magnesium ion and caused the β- and γ-phosphate groups of ATP to deviate from their catalytic orientations (**Fig. 2C-E**).

Remarkably, both FER mutants adopted an active conformation (**Fig. 3A-B**). Overall structural comparison of FER-KD^S525A^-ADP, FER-KD^K565R^-ADP and FER-KD^K565R^-ATP with the FER-KD-ADP complex gives Cα RMSD values of 0.230 Å, 0.280 Å and 0.187 Å, respectively (**Fig. 3A-B**). In particular, similar to the structure of FER-KD, the activation loop adopts an open conformation to allow substrate binding and phosphorylation; the C helix adopts an active state in which E581 forms an ion pair with K565 or R565 to contact the phosphate of the nucleotide; the regulatory spine is precisely formed; and the DFG motif belongs to the “BLAminus” type in the “DFGin” state, wherein the D679 of DFG points toward the active site to coordinate the phosphate of the nucleotide (**Fig. 3B**). Taken together, these data indicate that both the K565R and partially active S525A mutants adopted active forms without phosphorylation. Although no ATPase activity of FER-KD^K565R^ was detected (**Fig. 2F**), the ultrahigh concentration of FER-KD^K565R^ showed a weak autophosphorylation ability in an *in vitro* kinase assay (**Fig. S3B**). Our structural data and kinase assay were consistent with the genetic observation that FER-KD^K565R^ can partially replace FER-KD’s function (**29, 41**) since both FER-KD^K565R^ and FER-KD have similar structural forms (i.e., the active conformation).

**Figure 3.**
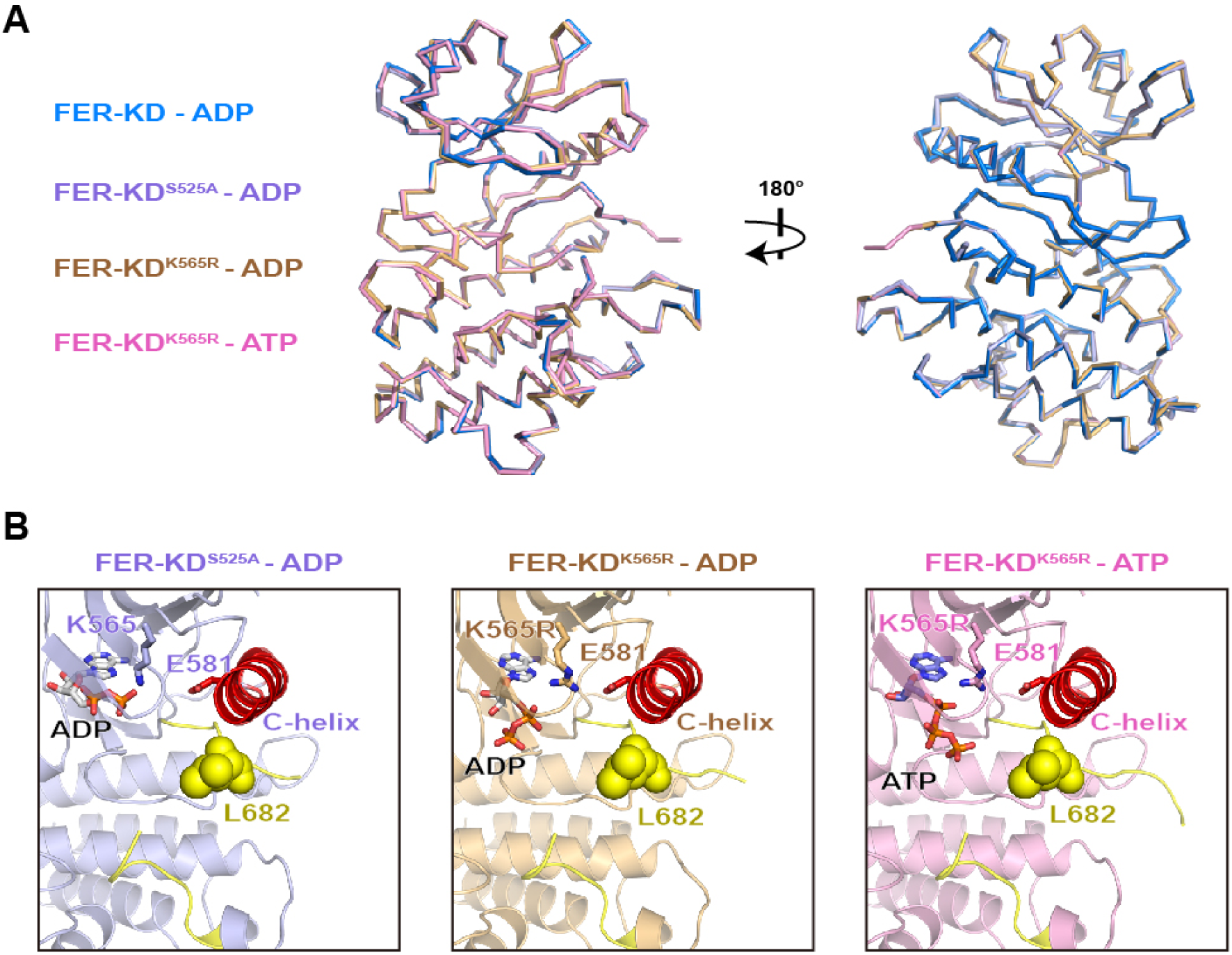
FER mutants fold into an active conformation identical to that of wild-type FER. **(A)** Structural comparison of the FER-KD-ADP complex (dark blue) with FER-KD^S525A^-ADP (light blue), FER-KD^K565R^-ADP (light orange) and FER-KD^K565R^-ATP (pink), which gives Cα RMSDs of 0.230 Å, 0.280 Å and 0.187 Å, respectively. Their conformations are almost the same. (B) Detailed comparison of the C-helix and activation segment of FER-KD^S525A^-ADP, FER-KD^K565R^-ADP, and FER-KD^K565R^-ATP. The activation segment is colored yellow. The C-terminal region is colored red. L682 is shown as yellow spheres. The residues K565/R565 and its conformation-activating residue E581 are shown as colored sticks.

### FER employs a *trans*-mode for phosphate transfer during autophosphorylation

The activity of protein kinases is regulated in several ways, with autophosphorylation activation being one that is the most commonly observed. To characterize the ATPase activity of FER-KD, we performed an assay using the NADH-coupled system (**50**) and determined the *Km* and *kcat* values to be 14.21 ± 2.166 μM and 0.09972 ± 0.004651 s^-1^, respectively. After conversion, *kcat*/*Km* is approximately equal to 0.007 μM^-1^ s^-1^. These results show that FER-KD could effectively hydrolyze ATP (**Fig. 4A**).

**Figure 4.**
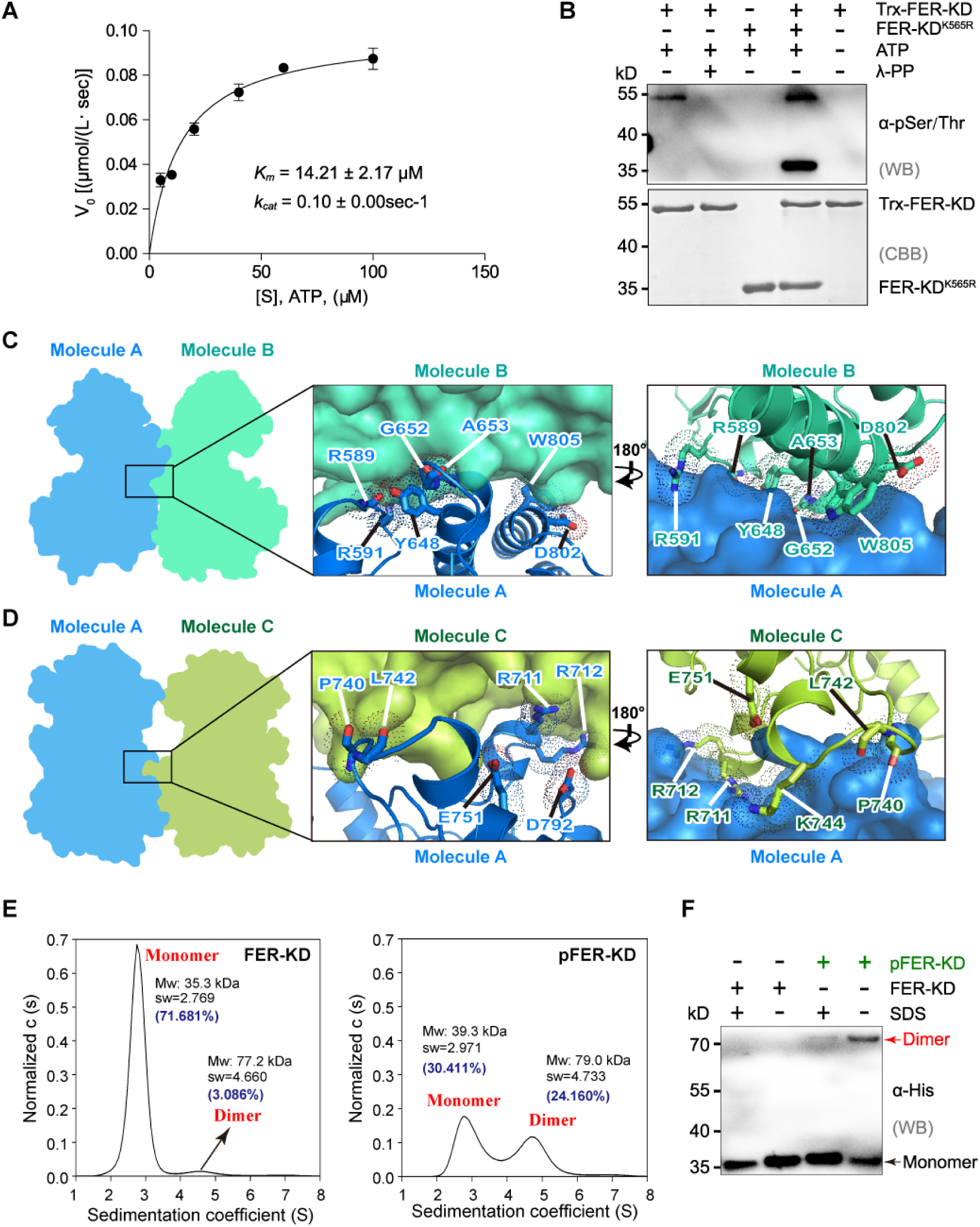
FER autophosphorylation is accomplished in a trans manner. **(A)** Measurement of kinetic parameters for FER-KD using the NADH-coupled method. (B) *Cis/trans*-autophosphorylation analyses of FER-KD by western blotting. The phosphorylation states of Trx-FER-KD and 6×His-FER^K565R^ were analyzed by using a general anti-pSer/Thr antibody. (C) Overview of the back-to-back dimer in the crystal and detailed view of the dimeric interface. Molecule B (green-cyan) is shown in surface representation in the first exploded view. Interfacial residues from molecule A (dark blue) are highlighted. In the second exploded view, molecule A is shown in surface representation, and interfacial residues from molecule B are highlighted. (D) Overview of the face-to-face dimer in the crystal and detailed view of the dimeric interface. Molecule C (green–yellow) is shown in surface representation in the first exploded view. Interfacial residues from molecule A are highlighted. In the second exploded view, molecule A is shown in surface representation, and interfacial residues from molecule C are highlighted. (E) Sedimentation velocity analytical ultracentrifugation of FER-KD and pFER-KD. (F) Native PAGE to analyze FER-KD and pFER-KD dimerization.

To further understand whether FER employs a *cis*-(intramolecular-) or *trans*-(intermolecular-) mode for phosphate transfer during autophosphorylation, we examined whether isolated Trx-FER-KD could phosphorylate purified FER-KD^K565R^ (see Methods for details). *In vitro* kinase assay results indicated that Trx-FER-KD could *trans*-phosphorylate the FER-KD^K565R^ mutant. In control experiments, the FER-KD^K565R^ mutant could not phosphorylate itself (**Fig. 4B**), and autophosphorylation of Trx-FER-KD could be effectively reversed by Lambda protein phosphatase (λ-PP), a broad-spectrum protein phosphatase (**51**) (**Fig. 4B**). Consistent with the notion that *trans*-autophosphorylation depends on the formation of kinase dimers (**52-54**), we analyzed and noted the existence of a back-to-back dimer and a face-to-face dimer in the crystal structure of FER-KD (**Fig. 4C-D**). Both types of dimers are connected primarily by polar contacts. There are 12 and 10 hydrogen bonds formed in the back-to-back and face-to-face dimers, respectively (**Fig. 4C-D**).

To investigate the oligomerization state of FER-KD in solution, we performed sedimentation velocity analytical ultracentrifugation (SV-AUC) on FER-KD and postphosphorylated FER-KD (pFER-KD, pretreated with 1 mM ATP-Mg^2+^ for 30 min) at concentrations of 19.7-22.5 μM. As shown in Figure 4E, FER-KD behaved mainly as monomers, with a trace number of dimers (3.086%), while phosphorylation of FER-KD induced the formation of a significant number of dimers (24.160%) (**Fig. 4E**). pFER-KD dimerization was also verified by native PAGE (**Fig. 4F**). These results demonstrate that autophosphorylation of FER RLK appears to dramatically stabilize the dimeric form of the kinase, which may significantly facilitate its full activation and signal transduction.

### The Ser/Thr/Tyr sites in the activation segment are essential for FER autophosphorylation

To further confirm the autophosphorylation activity of FER-KD, we set up an experiment with a time-gradient *in vitro* kinase assay. The results showed that Ser/Thr (S/T) of FER-KD could be phosphorylated at 1 min, and its phosphorylation level increased in a time-dependent manner and reached the apparent saturation state at approximately 30 min (**Fig. 5A**). In contrast, the phosphorylation of Tyr (Y) is much slower than that of S/T, as Tyr phosphorylation does not appear until 10 min (**Fig. 5A**). To determine specific phosphorylation sites in FER-KD, we performed a time-resolved (0 min, 1 min, 5 min, 10 min, 15 min and 30 min) autophosphorylation assay for FER-KD *in vitro* and identified the detailed autophosphorylation sites by LC– MS/MS (**Fig. 5B**). Consistent with our previous results (**Fig. S1A**), no phosphorylation could be detected at 0 min (**Fig. 5B**). We found five phosphorylation sites (T685/T688/T692/S695/T696) in the activation segment and three other phosphorylation sites (T533/T560/T656) at 1 min (**Fig. 5B**, **Fig. S4**), indicating that the eight phosphorylation sites may dictate the initiation of FER-KD autophosphorylation. Phosphorylation of Tyr does not appear until 10 min (**Fig. 5B**, **Fig. S2**), which is consistent with the *in vitro* kinase assay and suggests that the autophosphorylation of Tyr residues lags somewhat behind Ser/Thr phosphorylation. A total of 23 phosphorylation sites were identified *in vitro* at 30 min (**Table S3**). Among these, six sites were consistent with *in vivo* sites reported in published databases (**Fig. S5**) (**55-57**); one is S525, for which we already have the structure, and the other five are T688, T692, S695, T696 and S701 on the activation segment. Sequence alignment of the cytosolic domain from 17 members of the FER subfamily revealed that the relatively well-conserved phosphorylation sites were mainly located in the activation segment (residues 678-708) (**Fig. S2**).

**Figure 5.**
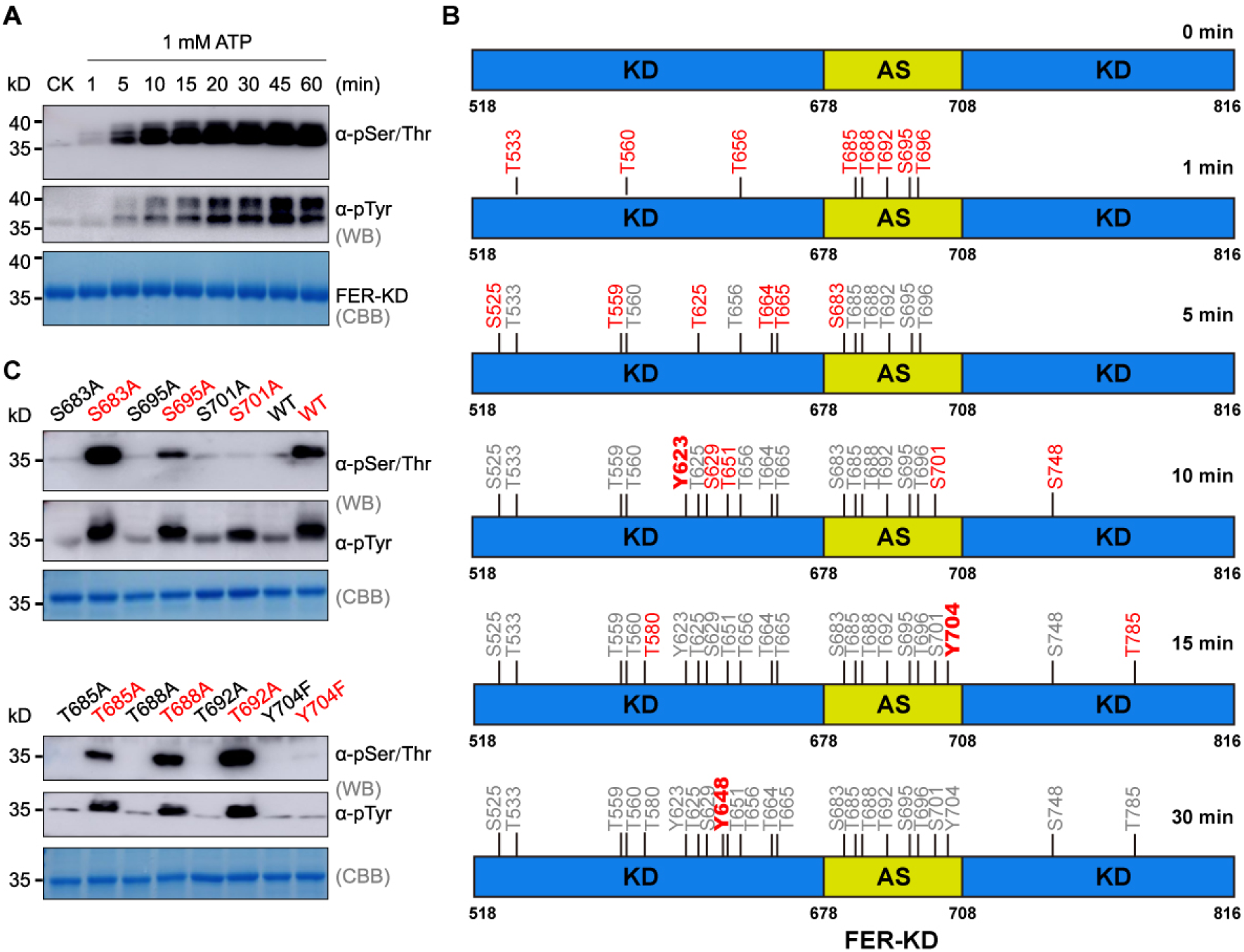
The significance of autophosphorylation on the activation segment of FER. **(A)** FER-KD time-gradient autophosphorylation assay. **(B)** MS identification of FER-KD autophosphorylation sites *in vitro* at different time points. The phosphorylation sites are numbered in the upper panel. Compared to the last point of time, newly emerging phosphorylation sites are shown in red. At one minute, 8 phosphorylation sites were detected. Near complete phosphorylation was found at 30 min. **(C)** Autophosphorylation activities of FER variants with a single mutation in the activation segment. Each dephosphorylated FER variant in the absence or presence of ATP and Mg^2+^ is indicated in black or red, respectively.

Phosphorylation of amino acid residues on the activation segment appears to be closely related to kinase activity and catalytic mechanism. Eight mutants were constructed with a single Ser/Thr-to-Ala or Tyr-to-Phe substitution to evaluate their autophosphorylation capacity by an *in vitro* kinase assay (**Fig. S3C**). The results showed that the single mutant of S683A, T685A, T688A, T692A and S695A retained autophosphorylation capacity, as Ser/Thr/Tyr phosphorylation was found in the *in vitro* kinase assay (**Fig. 5C**), but the mutant of Y704F lost autophosphorylation capacity (**Fig. 5C**). We further found that the autophosphorylation capacity the mutant of T696A was significantly reduced and the double mutant of S695A/T696A lost the autophosphorylation capacity **Fig. S6A**). Interestingly, the mutant of S701A lost Ser/Thr autophosphorylation activity but retained Tyr autophosphorylation activity (**Fig. 5C**).

### Tyr phosphorylation is essential for FER-KD to initiate substrate phosphorylation

To further investigate the effect of autophosphorylation on FER-KD, we prepared pFER-KD based on the results of the time-gradient autophosphorylation assay. This phosphorylation could be effectively reversed by λ-PP (Fig. S6B). Furthermore, we used the previously reported FER phosphorylation substrate glycine-rich RNA-binding protein 7 (GRP7) (**58**) as the substrate to verify the transphosphorylation activity of FER-KD.

With an increased amount of ATP, FER-KD would first turn on the autophosphorylation reaction and then start the phosphorylation of GRP7 (transphosphorylation reaction) when its autophosphorylation reached a steady level (1 mM ATP for 30 min) (**Fig. 6A**). With 1 mM ATP, the phosphorylation of GRP7 by FER-KD showed a similar time-gradient dependence. The ability of FER-KD to phosphorylate GRP7 was relatively weak in the first 10 minutes; however, FER-KD showed very strong transphosphorylation of GRP7 at 30 minutes (**Fig. 6B)**. In sharp contrast, pFER-KD immediately initiated the reaction to transphosphorylate GRP7 (**Fig. 6B)**. Through careful analysis of time-gradient phosphorylation and autophosphorylation sites by LC–MS/MS (**Fig. 5B**), we observed that the phosphorylation level of Ser/Thr increased rapidly and concluded within 15 min (**Fig. 6B)**, while the phosphorylation level of Tyr increased slowly. Interestingly, the activation of FER-KD substrate GRP7 phosphorylation was time-consistent with the autophosphorylation of Tyr (**Fig. 6C**). This indicates that if the pFER-KD protein is well phosphorylated with basal Tyr phosphorylation (**Fig. 6B**), it could ensure the subsequent substrate GRP7 phosphorylation, leading to the onset of downstream cellular processes (**Fig. 6B**). This hypothesis is further supported by the observation that three mutants (Y623F, Y704F and Y709F) lost the ability to phosphorylate the substrate GRP7 (**Fig. 6D**), and that although Y648F retains the ability of autophosphorylation, its ability to phosphorylate GRP7 is greatly reduced (**Fig. 6D**). Our data indicate that during signal initiation, FER-KD is likely to undergo Ser/Thr autophosphorylation first, then Tyr autophosphorylation, and finally initiate substrate phosphorylation (**Fig. 6A-C**). These results indicate that Tyr phosphorylation gates substrate phosphorylation of FER.

**Figure 6.**
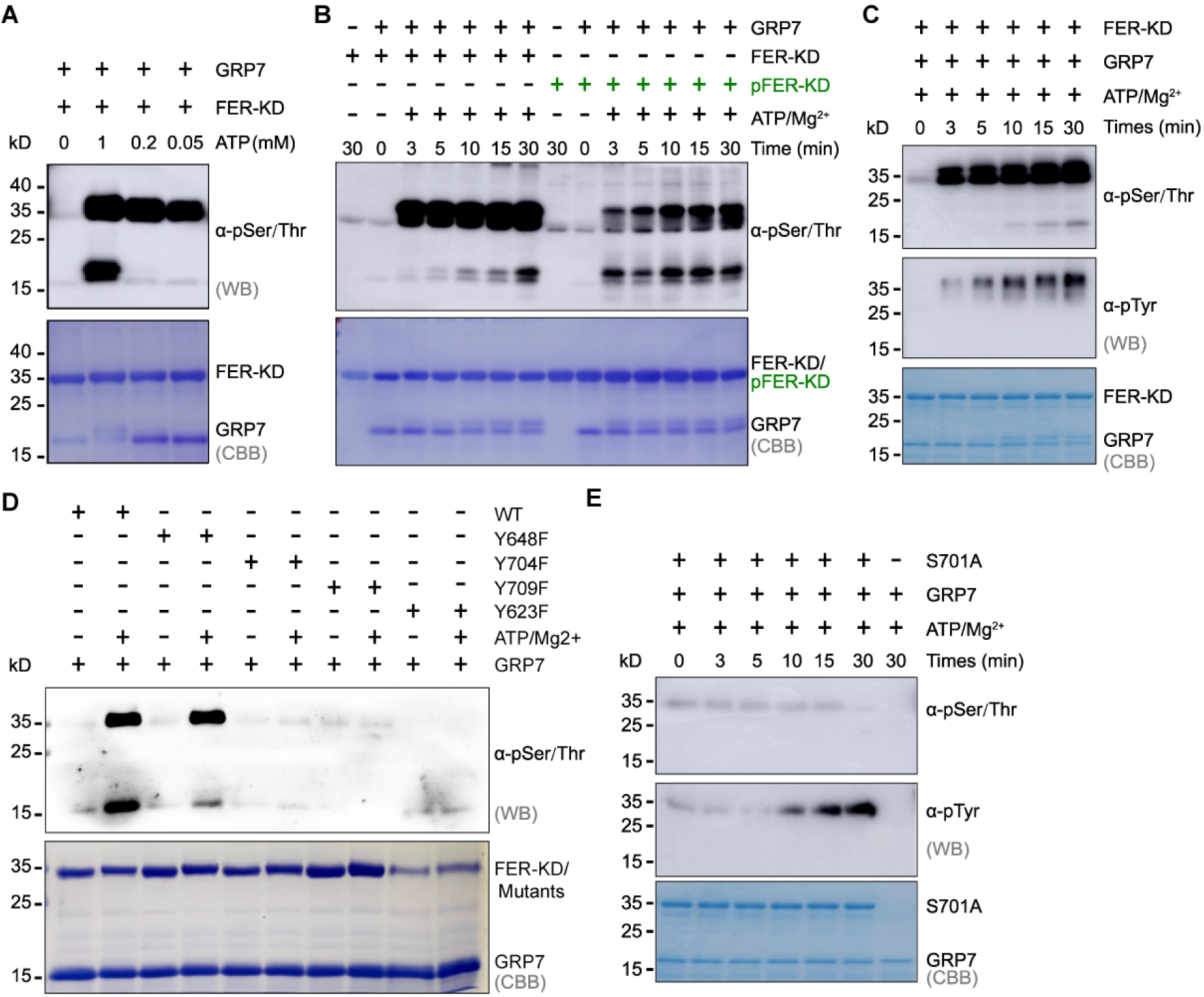
Tyr phosphorylation gates substrate phosphorylation of FER. **(A)** Substrate reactions at different ATP concentrations. FER-KD was reacted with GRP7 at different concentrations of ATP for 30 minutes at 25°C. (B) Time-resolved phosphorylation of GRP7 by dephosphorylated and phosphorylated FER-KD. The same molar mass of FER-KD/pFER-KD was reacted with GRP7 for the time-gradient at 25°C. Phosphorylated pFER-KD samples are shown in green. (C) Transphosphorylation activities assay of FER-KD to the GRP7 substrate. (D) Transphosphorylation activities assay of FER-KD^Y623F^, FER-KD^Y648F^, FER-KD^Y704F^ and FER-KD^Y709F^ to the GRP7 substrate. (E) Transphosphorylation activities assay of FER-KD^S701A^ to the GRP7 substrate.

The autophosphorylation assay mentioned in the previous section identifies that the S701A mutant exhibits compromised Ser/Thr autophosphorylation activity yet promoted Tyr autophosphorylation activity (**Fig. 5C**). We used this distinctive characteristic of S701A to further confirm its effect on FER substrate transphosphorylation activity. This mutant revealed an undetectable level of S/T autophosphorylation and an elevated level of Tyr autophosphorylation, as expected (**Fig. 6E**). Although the autophosphorylation of Tyr increased, the substrate transphosphorylation activity of FER-KD was still scarce (**Fig. 6E**). These results suggest that autophosphorylation of Tyr is necessary but not sufficient to promote FER-regulated substrate phosphorylation. It appears that both Ser/Thr autophosphorylation and Tyr autophosphorylation need to be satisfied prior to the initiation of FER-KD substrate phosphorylation activity.

## CONCLUSION

Our work reports, for the first time, a phosphorylation-independent active state of RLKs in plants. Although RLK FER utilizes a phosphorylation-independent active state, it switches and ensures subsequent substrate phosphorylation in a delayed Tyr phosphorylation-gated manner, providing a paradigm to study the mechanisms of the early stages during RLK signaling initiation.

Although this is the first report of a phosphorylation-independent active conformation in plants, its counterpart in animal systems was not unprecedented. For example, both cyclin-dependent kinase 2 (CDK2) and epidermal growth factor receptor (EGFR) are activated by intermolecular interactions instead of phosphorylation (**59, 60**). CDK2 forms an inactive conformation in isolation and switches to an active conformation upon cyclin A binding but not through protein phosphorylation, while EGFR forms an active conformation by homo or heterodimerization. The phosphorylation-independent active state of FER appears to be functionally relevant. For example, the kinase-dead FER^K565R^ protein was able to complement ovule fertilization (**41**) and PT reception defects in the *fer-4* mutant (**27**) and root mechanosensitive defects in the *fer-2* and *fer-4* mutants (**61**). FER in the active state may likely facilitate, in part, recruitment or activation of specific downstream substrates for signal transduction.

While the substrate phosphorylation activity of FER-KD is not activated by the change in active site conformation, our evidence suggests that it is rather activated by Tyr autophosphorylation in a dose-dependent manner. Delayed Tyr phosphorylation-gated substrate phosphorylation may serve as an essential mechanism to avoid the noise response caused by RLKs at the PM. In other words, short-term and/or low-dose ligand stimulation without Tyr phosphorylation will not lead to the onset of a multitude of downstream cellular processes. Thus, Tyr phosphorylation functions as a biological noise filter in RLKs.

Furthermore, an *in vitro* kinase assay revealed that high-concentration “kinase-dead” FER-KD^K565R^ caused a weak autophosphorylation ability (**Fig. S3B**). This is consistent with our previous work showing that FER-KD^K565R^ has a weak ability to trigger eIF4E phosphorylation *in vivo* (**36**). This can partially explain why FER^K565R^ can rescue a limited set of cellular processes (e.g., pollen tube growth regulation) of the FER mutant. Pseudokinases account for ∼20% of all RLKs that exist in plants and are more likely to adopt a phosphorylation-independent active state mechanism by definition (**21**). More studies focusing on RLK phosphorylation-independent active state mechanisms may also facilitate understanding the essential roles of pseudokinases in distinct biological processes in plants.

## MATERIALS AND METHODS

### Protein expression and purification

FER-KD (residues 518-816) and all point mutations were cloned into the pRSF-Duet vector with an N-terminal 6×His tag. Trx-FER-KD was cloned into pET32a with a Trx tag, a 6×His tag and an S-tag in tandem at the N-terminus, in which thioredoxin (Trx) is an approximately 12 kDa soluble *E. coli* protein that is used as a tag. FER-KD, Trx-FER-KD and the FER-KD^S525A^ mutant were coexpressed with the Arabidopsis phosphatase ABI2, which was cloned into the pGEX-4T-1 vector (**44**). Other mutants were coexpressed with λ-phosphatase (λ-PP), which was cloned into the pGEX-6P-1 vector. GRP7 was cloned into the pGEX-6P-1 vector. All proteins were expressed in the same *E. coli* strain, BL21(DE3). Protein expression was induced at 289 K for 18 h with 1 mM isopropyl-β-D-thiogalactopyranoside (IPTG) after the OD_600_ reached 0.6. The cells were harvested by centrifugation at 5000 rpm for 15 min, resuspended in phosphate-buffered saline (PBS, pH 7.5) containing 10 mM imidazole, and crushed by using a low-temperature ultrahigh-pressure cell disrupter (JNBIO, Guangzhou, China). The soluble proteins were separated from cell debris by high-speed centrifugation (12,000 rpm for 1 h), and 6×His-tagged proteins were loaded onto a Ni^2+^ affinity column (Smart Life Sciences, SA004250) and washed with 100 ml of washing buffer (PBS containing 20-100 mM imidazole). The target protein was eluted from the column with PBS containing 300 mM imidazole, and GRP7 proteins were hung on glutathione beads (Smart Life Sciences, SA008100), washed with 60 ml of washing buffer (PBS 7.5), digested overnight with prescission protease or not and eluted with PBS or PBS containing 10 mM GSH. All proteins were exchanged into Tris buffer (20 mM Tris-HCl, 25 mM NaCl; pH 7.5) by using centrifugal concentrators (30 kDa, Millipore, UFC901096) and concentrated to approximately 6 mg/ml for storage (193 K).

To assemble the FER-KD-ATP/ADP complex, 3 mg/ml purified FER-KD, FER-KD^K565R^ and FER-KD^S525A^ were incubated with 10 mM ATP/ADP at 277 K for 12 h.

### Protein crystallization and data collection

Crystallization of the FER-ATP/ADP complex was accomplished by using the hanging drop vapor diffusion method by mixing 1 μl of protein complex with 1 μl of reservoir solution in 24-well plates at 291 K (Hampton Research, Aliso Viejo, CA). We successfully obtained crystals with high diffraction quality for the following complexes: FER-KD-ADP, FER-KD^K565R^-ATP, FER-KD^K565R^-ADP and FER-KD^S525A^-ADP. The best crystals were directly soaked in cryoprotectant solution (reservoir solution supplemented with 30% (v/v) glycerol) and flash frozen in liquid nitrogen at 100 K. X-ray diffraction data were collected on beamline BL17U1 at the Shanghai Synchrotron Radiation Facility (SSRF) using an ADSC Q315 CCD detector. Datasets were processed and scaled by using DIALS (**62**). Data collection statistics are summarized in Supplementary Table 1.

### Structure determination and refinement

The crystal structure for each of the four complexes (FER-KD-ADP, FER-KD^K565R^-ATP, FER-KD^K565R^-ADP and FER-KD^S525A^-ADP) was determined by molecular replacement with the atomic coordinates of the Pto kinase in the Pto-AvrPto complex (PDB entry 2QKW) as the initial search model. The atomic models were manually built in COOT (**63**) and refined in PHENIX (**64**). Analysis with MOLPROBITY within the PHENIX suite (**65**) suggested correct stereochemistry, with the fewest outliers in the Ramachandran plot. The refinement statistics are summarized in Supplementary Table S1. The atomic coordinates and structure data (PDB codes 7XDY for FER-KD-ADP, 7XDX for FER-KD^S525A^-ADP, 7XDW for FER-KD^K565R^-ADP and 7XDV FER-KD^K565R^-ATP) have been deposited in the Protein Data Bank. All structural figures were prepared with PyMOL (The PyMOL Molecular Graphics System, Schrödinger, LLC).

### NADH-coupled enzyme system

The NADH system involves the reaction of the ADP released by the phosphorylation process with enolpyruvate phosphate (PEP) catalyzed by pyruvate kinase (PK) to form pyruvate and ATP, followed by the reaction of pyruvate with NADH catalyzed by lactate dehydrogenase (LDH) to form NAD and lactate. Thus, the phosphorylation process consumes one molecule of NADH for each molecule of ADP produced so that the decrease in the light absorption of NADH at 340 nm (ε340 = 6,220 cm-1 M-1) can be monitored by using a UV spectrophotometer. Another feature of this system is that once ADP is produced, it is coupled to the enzymatic reaction to produce ATP, and thus, the total amount of ATP in the reaction system is not depleted. Protein viability assays were performed at 25°C in a chamber containing 20 mM Tris (pH 7.5), 10 mM MgCl2, 200 μM NADH, 1 mM PEP, 1 mM ATP, 20 units/mL LDH, 15 units/mL PK and different concentrations of substrate in a 200 μL viability buffer system. Each assay was repeated three times, and similar results were obtained.

### *In vitro trans*- or *cis*-autophosphorylation assays

For *in vitro trans/cis*-autophosphorylation assays, purified Trx-FER-KD and FER-KD^K565R^ proteins were treated with 1 μl of λ-phosphatase (λ-PP) (400,000 units/ml, New England Biolabs. P0753S) and 1 mM MnCl2 for 1-2 h at room temperature. The proteins were then buffer-exchanged into 50 mM HEPES (pH 7.5) using an ultrafiltration device (Millipore, UFC901096). The Trx-FER-KD and FER-KD^K565R^ proteins (1 μM) were coincubated at room temperature for 30 min in 50 μl of assay buffer containing 1 mM ATP and 5 mM Mg^2+^. Proteins were resolved by SDS–PAGE and either stained with Coomassie blue G250 or transferred to a nitrocellulose membrane (PALL, P-N66485) for protein immunoblotting using anti-phosphothreonine (Abcam, ab218195), anti-phosphoserine (Abcam, ab9332) and anti-phosphotyrosine (Abcam, ab179530) antibodies.

### *In vitro* kinase assay

For *in vitro* kinase assays, GRP7, FER-KD and FER-KD mutant proteins that coexpressed with λ-PP were purified. The FER-KD or FER-KD mutant proteins (1 μM) were coincubated with GRP7 (1.5 μM) at room temperature for 30 min or the time-gradient in 50 μl of assay kinase buffer containing 1 mM ATP and 10 mM Mg^2+^. Proteins were resolved by SDS–PAGE and either stained with Coomassie blue G250 or transferred to a nitrocellulose membrane (PALL, P-N66485) for protein immunoblotting using anti-phosphothreonine (Abcam, ab218195), anti-phosphoserine (Abcam, ab9332) and anti-phosphotyrosine (Abcam, ab179530) antibodies.

### Sequence alignment

The amino acid sequences of FER subfamily proteins from *Arabidopsis thaliana* were obtained from The Arabidopsis Information Resource (TAIR) site. The sequences were subjected to multiple-sequence alignment (MSA) using the T-Coffee server (**66**), and the MSA results were sent to figure production by ESPript version 3.0 (**67**).

### Analytical ultracentrifugation (AUC)

To assess the solution state of FER-KD and pFER-KD (pretreated with ATP-Mg^2+^ for 30 min), sedimentation velocity analytical ultracentrifugation (SV-AUC) experiments were performed using an XL-I analytical ultracentrifuge (Beckman Coulter) at 20°C. FER-KD/pFER-KD was prepared in a buffer containing 20 mM Tris pH 7.5, 150 mM NaCl, and the protein concentration was controlled at 0.4-0.8 mg/ml. Each 400 μl sample will be centrifuged at 50,000 rpm for 8 h in an An50Ti rotor using 12 mm double-sector aluminum centerpieces. Samples are recorded every few minutes to form data. Data were then analyzed using SEDFIT 11.7 (**68**), GUSSI and SEDPHAT software. Theoretical settlement data were calculated using HydroPro 7C (**69**) with a hydrated radius of 3.1 Å for the atomic elements.

## ACKNOWLEDGMENTS

We cordially thank the staffs from beamline BL17U1 of Shanghai Synchrotron Radiation Facility (SSRF) for assistance during data collection. We also thank Dongdong Li (Tsinghua University) for assistance with AUC data analysis. **Funding:** This work was supported by grants from the National Natural Science Foundation of China (32160064, 31871396, 31571444, 32000916), the State Key Laboratory for Conservation and Utilization of Subtropical Agro-bioresources (SKLCUSA-a201806), China Postdoctoral Science Foundation (2019M662764), the Guangxi Natural Science Foundation (2020GXNSFFA297007), and the Guangxi Key Laboratory for Sugarcane Biology (GXKLSCB-20190304).

## AUTHOR CONTRIBUTIONS

Z.H.M and F.Y. conceived the project and designed the research; Y.Q.K, J.C, H.C, Y.N.S, L.F.W, and Y.J.Y performed the research; Y.Q.K, H.Z collected data; Z.H.M, F.Y, H.P.Z, Y.Q.K, and J.C analyzed data and wrote the paper; all authors reviewed and approved the manuscript for publication.

## Supplemental Data

**Figure S1.**
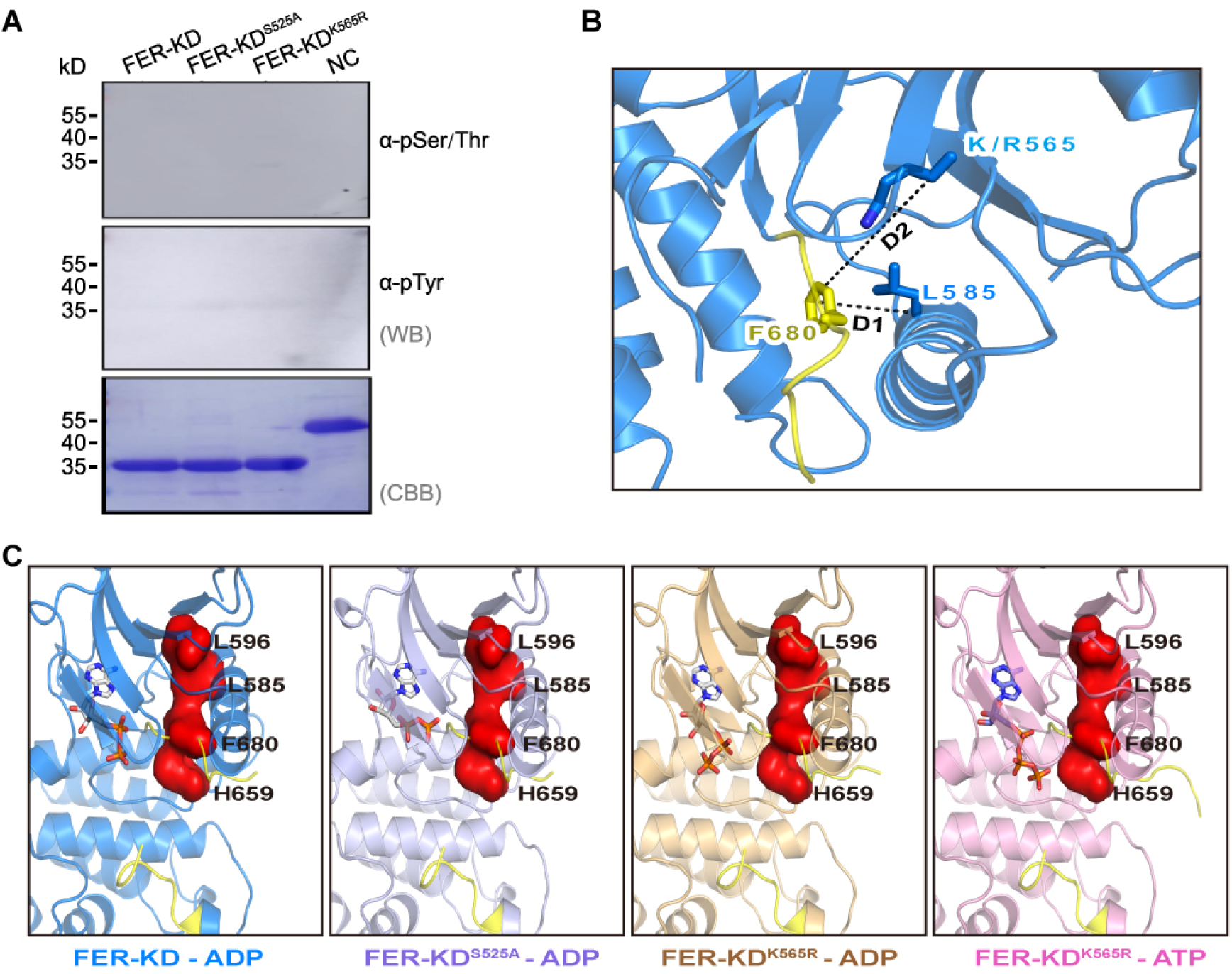
The purification and structures of various FER forms. **(A)** Validation of the phosphorylation state of the three crystallized proteins. **(B)** D1 and D2 indicate the distances between F680 and L585/K(R)565 in the DFG motif. **(C)** Conformations of the regulatory spine in different FER complexes. The spine is shown as a red surface.

**Figure S2.**
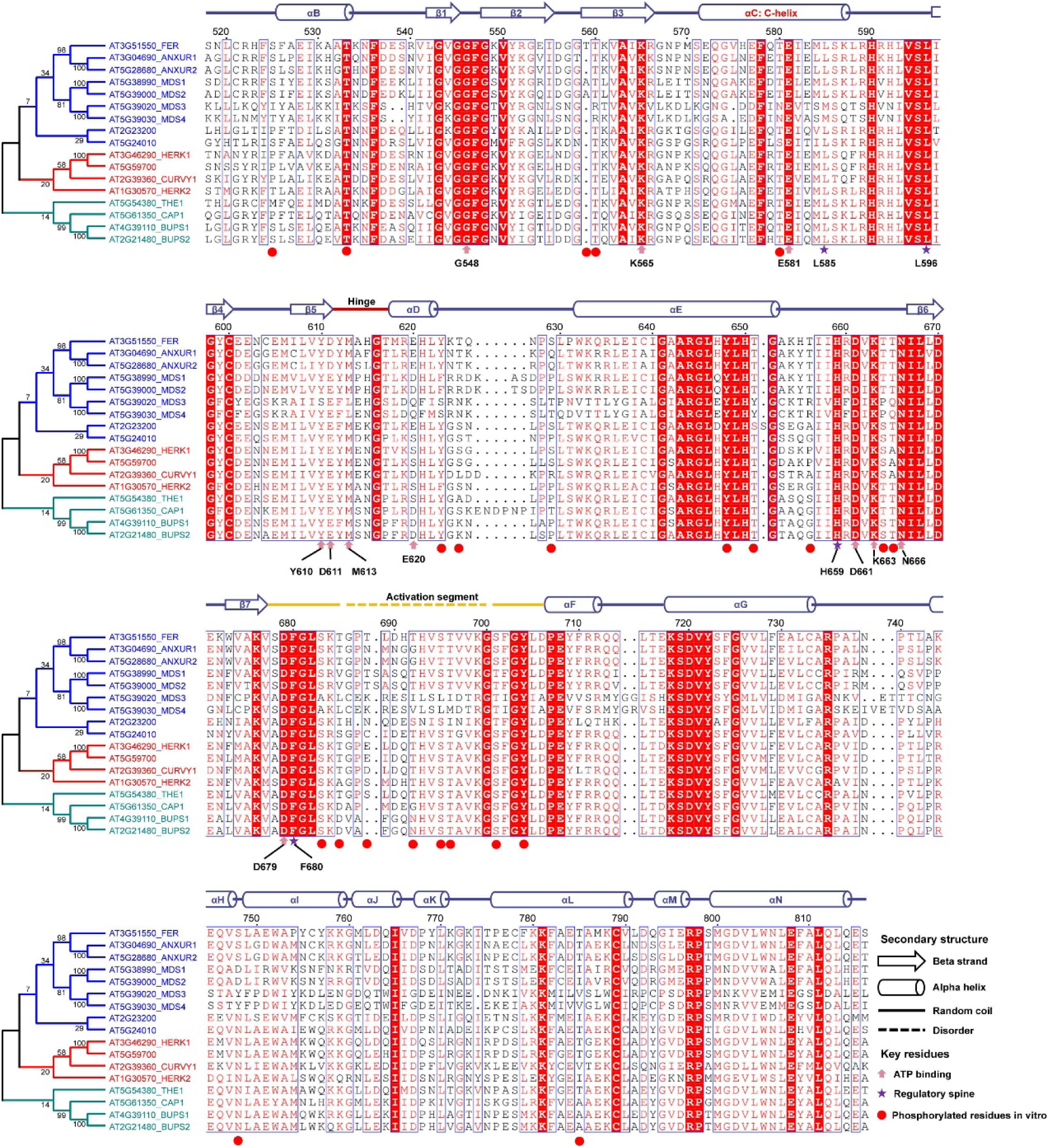
Structural elements, autophosphorylation sites and conservation of FER subfamily proteins. Sequence alignment of FER with sixteen representative FER subfamily proteins. Conserved amino acids are highlighted in the blue box. Secondary structural elements of FER are shown above the alignment. The autophosphorylation sites of FER are shown by solid red circles. The ATP-binding sites of FER are shown by solid pink arrows. The regulatory spine of FER is shown by a solid purple star.

**Figure S3.**
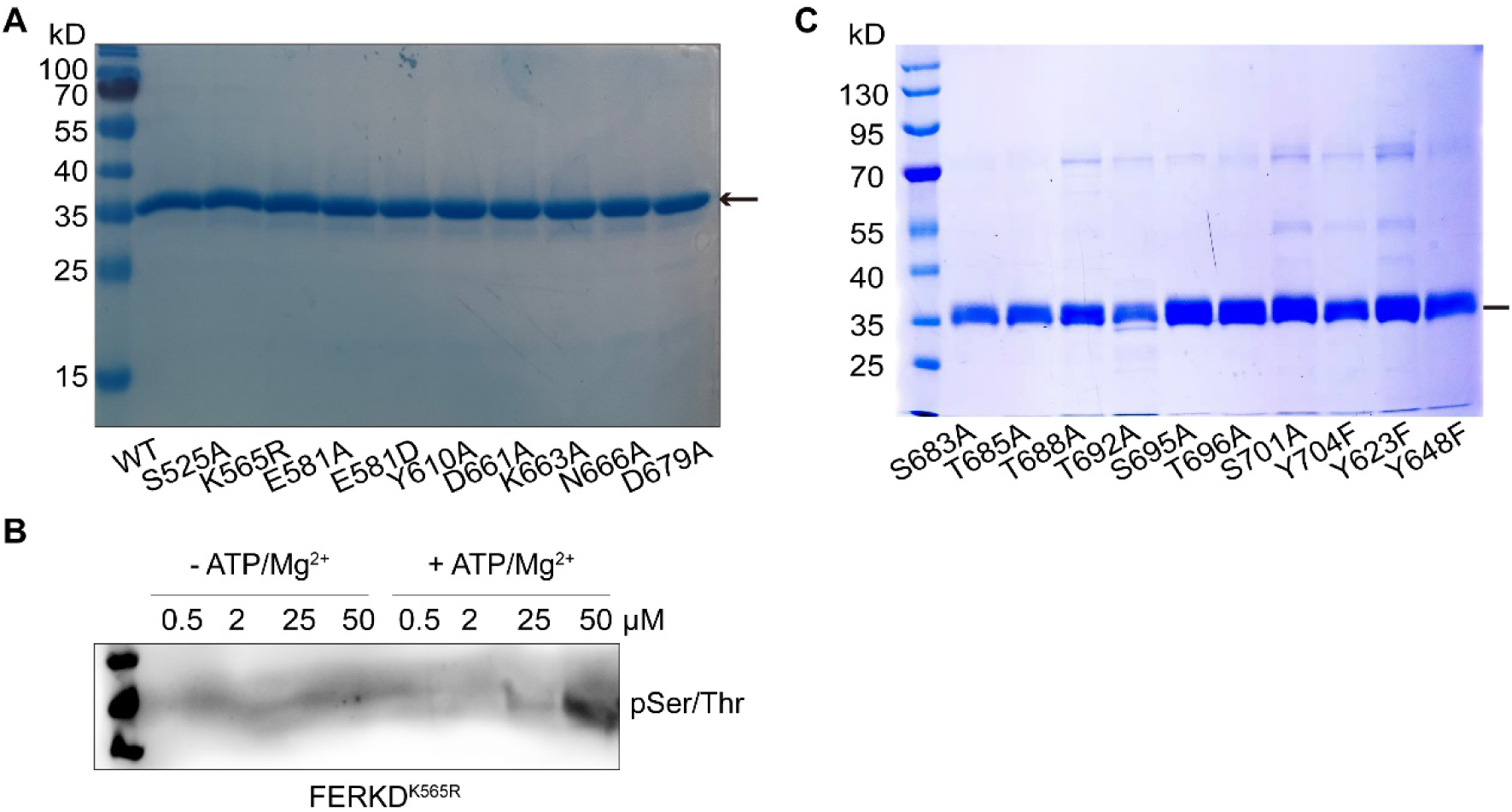
SDS–PAGE analysis of purified FER and the autophosphorylation of FER-KD^K565R^. **(A)** SDS–PAGE analysis of FER mutants affecting nucleotide binding. **(B)** Validation of autophosphorylation of FER-KD^K565R^ at ultrahigh concentrations. **(C)** SDS–PAGE analysis of the purified FER-KD mutants associated with the activation segment.

**Figure S4.**
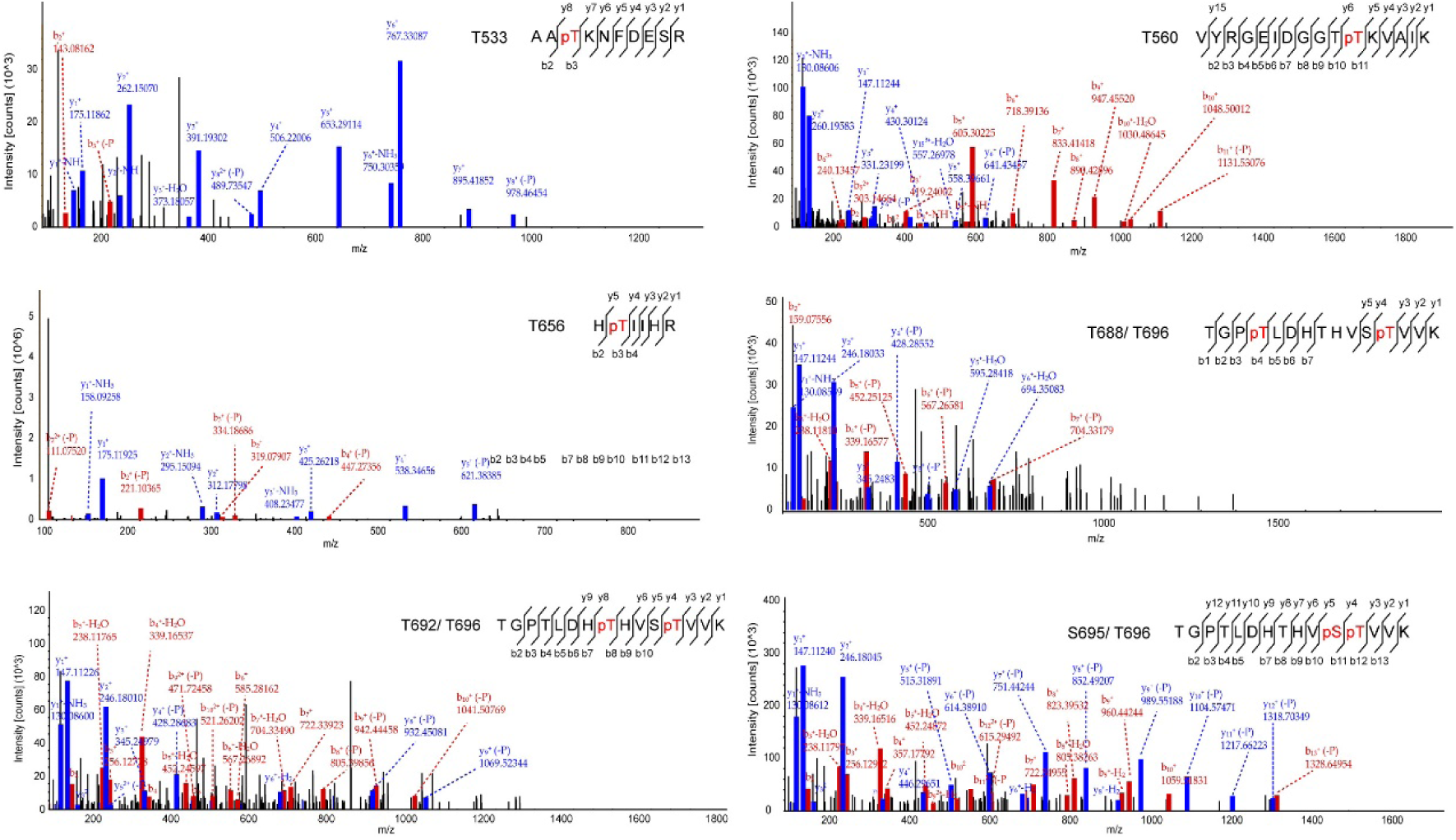
FER phosphorylation site identification. LC–MS/MS analysis was performed using an Easy-nanoLC-1000 coupled to an Orbitrap Elite mass spectrometer, identifying FER-KD autophosphorylation sites. The mass spectra of 8 phosphorylated FER-KD peptides at 1 min are shown.

**Figure S5.**
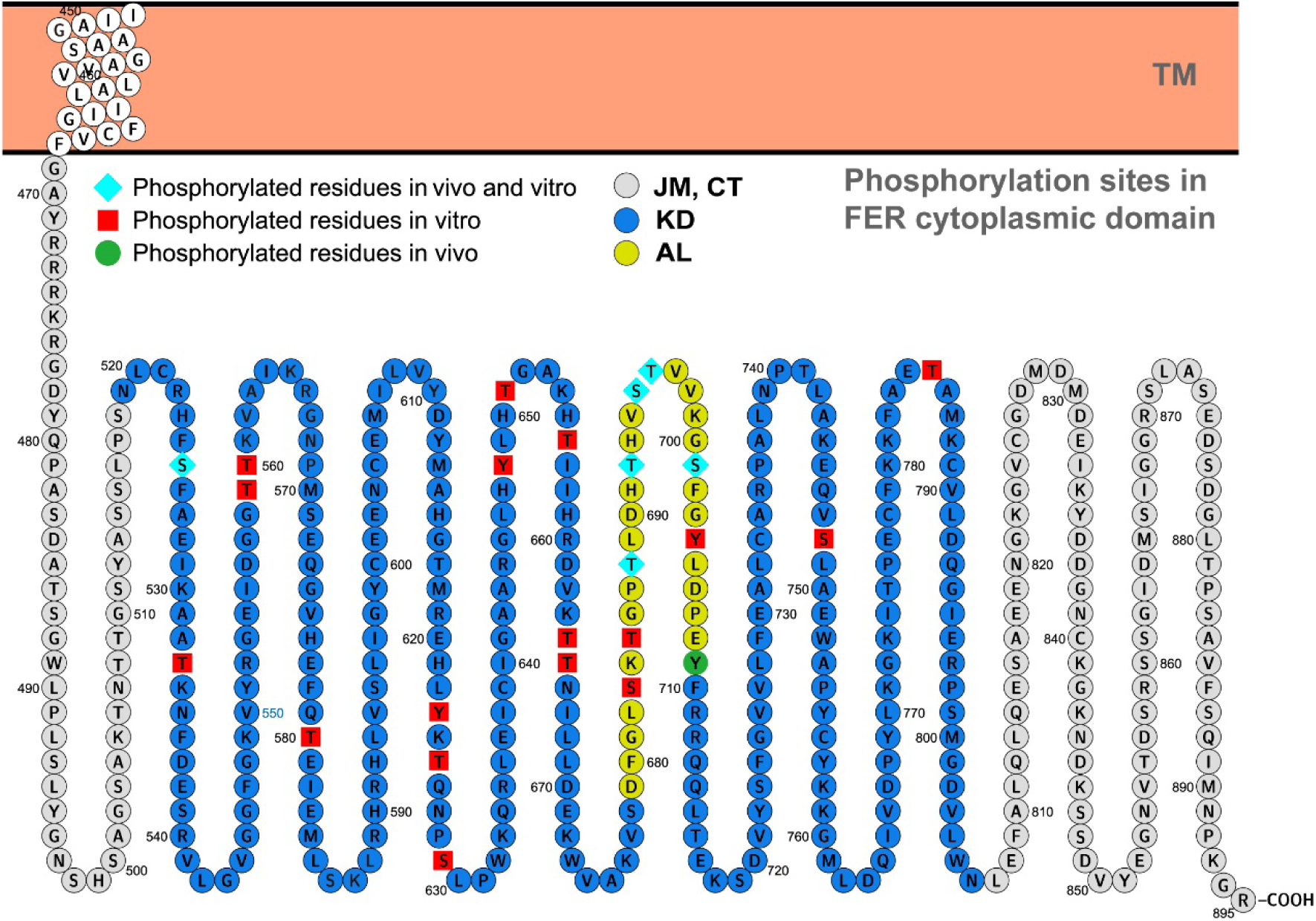
Visualization of FER phosphorylation sites. Visualization of MS/MS identification of FER autophosphorylation sites *in vivo* and the phosphorylation sites identified *in vitro*.

**Figure S6.**
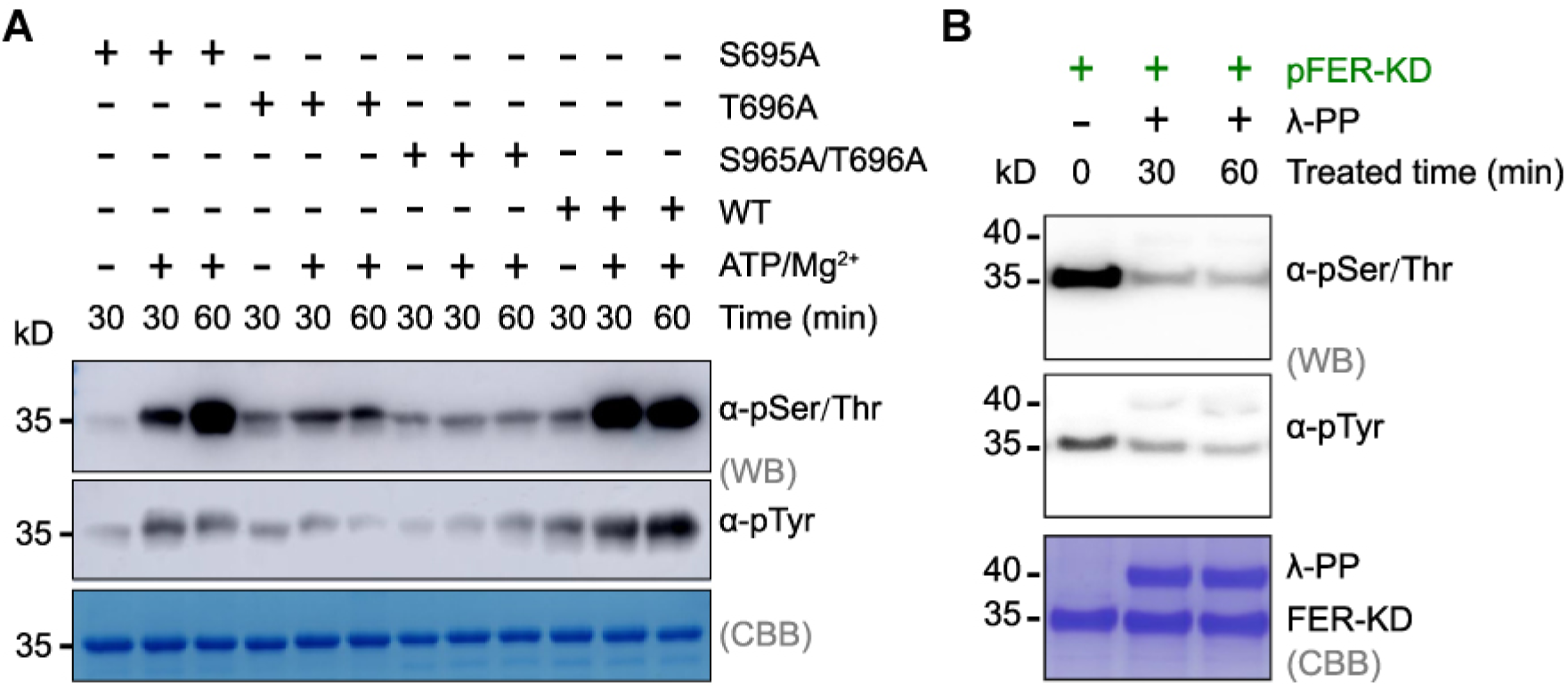
Autophosphorylation activities assay and preparation of FER-KD phosphorylated samples (pFER-KD) **(A)** Autophosphorylation activities assay the single mutant of S695A, T696A and double mutant of S695A/T696A in the activation segment. **(B)** Preparation of pFER-KD samples. Purified FER-KD protein after 1 mM ATP, 25 ℃, and 30 min reaction and buffer-exchanged into 20 mM Tris (pH 7.5) using centrifugal concentrators (Millipore, UFC901096). Phosphorylated pFER-KD samples are shown in green and could be effectively reversed by λ-PP.

**Table S1.** Analysis of kinase structures depending on the spatial position of the DFG-Phe side chain. According to the scatterplot criterion, FER belongs to DFGin.

|  | Chain | D1 | D2 | Conclusion |
| --- | --- | --- | --- | --- |
| FER-KD -ADP | A | 6.8 | 15.8 | DFGin |
|  | B | 6.4 | 16 | DFGin |
| FER-KD<sup>S525A</sup> -ADP | A | 6.9 | 15.5 | DFGin |
|  | B | 6.8 | 15.4 | DFGin |
| FER-KD<sup>K565R</sup> -ADP | A | 6.7 | 15.7 | DFGin |
|  | B | 6.8 | 15.6 | DFGin |
| FER-KD<sup>K565R</sup> -ATP | A | 6.7 | 15.6 | DFGin |
|  | B | 6.7 | 15.7 | DFGin |

**Table S2.** A finer analysis based on the dihedral angles required to place the Phe side chain. X-DFG (residue before the DFG motif), DFG-Asp, DFG-Phe and DFG-Gly. The ramachandran regions are marked A (alpha), B (beta), L (left), and E (epsilon). The DFG-Phe χ1 rotamer (minus = -60°, plus = +60°, and trans = 180°). According to the Measure standard, FER belongs to BLAminus.

|  | Chain | Ser-DFG |  | DFG-Asp |  | DFG-Phe |  | DFG-Phe | Conclusion |
| --- | --- | --- | --- | --- | --- | --- | --- | --- | --- |
| | | PHI | PSI | PHI | PSI | PHI | PSI | $\chi_1$ rotamer | |
| FER-KD -ADP | A | -141.8 | -171.9 | 66.9 | 59.9 | -83.4 | 11.2 | -75.1 | BLAminus |
|  | B | -144.2 | -178.1 | 65.9 | 74.2 | -94.9. | 27.1 | -68.0 | BLAminus |
| FER-KD<sup>S525A</sup> _<br>ADP | A | -133.4 | -178.7 | 67.5 | 77.5 | -97.3. | 15.8 | -74.0 | BLAminus |
|  | B | -136.8 | -177.1 | 63.7 | 79. | -96.7. | 14.6 | -74.1 | BLAminus |
| FER-KD<sup>K565R</sup> _<br>ADP | A | -137.4 | -174.6 | 58.4 | 60.6 | -84.7 | 27.0 | -80.0 | BLAminus |
|  | B | -133.7 | -171.6 | 65.1 | 59.0 | -87.1 | 21.9 | -79.7 | BLAminus |
| FER-KD<sup>K565R</sup> _<br>ATP | A | -134.3 | 177.5 | 68.4 | 79.2 | -98.6 | 11.5 | -77.4 | BLAminus |
|  | B | -133.7 | 179.3 | 66.8 | 78.3 | -97.8 | 11.6 | -80.1 | BLAminus |

**Table S3.**
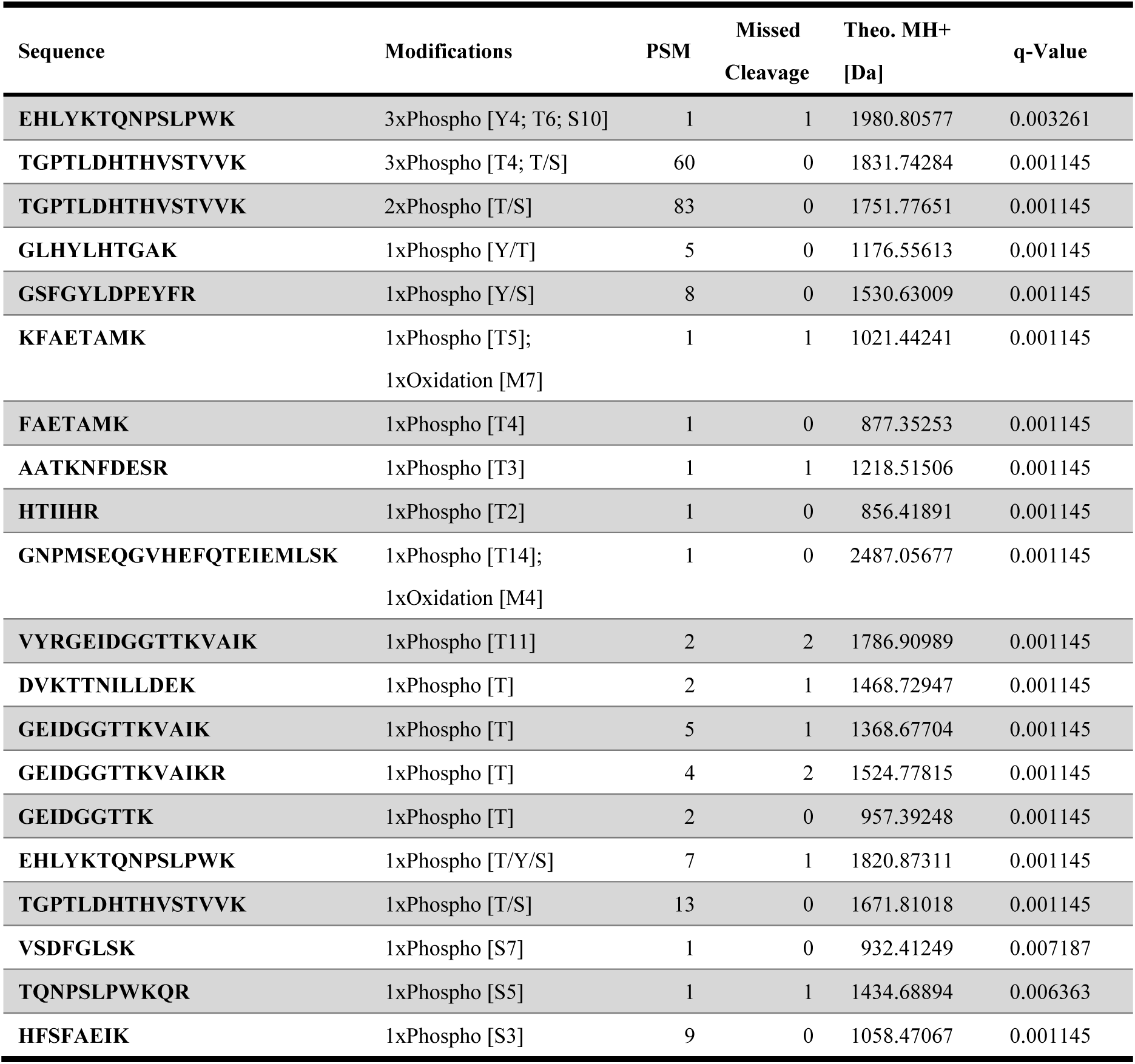
Mass spectrum of 23 phosphorylated FER-KD peptides at 30 min.

## Notes

### Competing Interest Statement

The authors have declared no competing interest.

